# Brain-wide single-neuron bases of working memory for sounds in humans

**DOI:** 10.1101/2025.11.10.687666

**Authors:** Joel I. Berger, Alexander J. Billig, Ryan M Calmus, Christopher K. Kovach, Ariane E. Rhone, Christopher M. Garcia, Sukhbinder Kumar, Hiroto Kawasaki, Matthew A. Howard, Timothy D. Griffiths

## Abstract

In order to understand the constantly changing acoustic world our brains must maintain elements of auditory scenes in memory. The neural mechanisms for this fundamental process remain unclear. Here, we report human intracranial recordings of 1269 single neurons while participants performed a non-verbal auditory working memory task that required adjustment of a tone frequency to match a target. We found neurons within hippocampus, insula and cingulate cortex, for which firing rates were modulated at different phases of the task, particularly during maintenance and adjustment. For the majority of the neurons modulated during maintenance there was a striking suppression of activity rather than increased activity. Across the entire neuronal population, state-space analysis demonstrated attractor-like states corresponding to different task phases. Behaviorally, more neurons were modulated at the beginning of the maintenance phase in trials when participants performed better. State-space analysis was consistent with greater attractor-like activity during the adjustment phase supporting better performance. These data support the existence of a widely distributed neural code for auditory working memory that determines related behavior based on attractor-like states.

## Main

To understand the acoustic world, animals and humans need to perceive current sound input and recall recent inputs. Based on human data utilizing magnetoencephalography, functional magnetic resonance imaging (fMRI) and local field potential (LFP) recordings, recall of sounds over seconds –working memory for sound – is associated with ensemble neuronal activity changes in human sensory cortex, but also in a network including frontal cortex and the hippocampus e.g. *[1-3]*.

Recent visual studies have examined human single-neuron activity while participants perform working memory tasks *[4-9]*, highlighting the involvement of brain regions including sensory cortex, but also hippocampus, amygdala and prefrontal cortex. In the auditory domain, single-neuron activity has only been examined during phonological working memory, showing a relationship between persistent activity and working memory load in hippocampus *[10]*. In the present study we report the first human single neuron recordings across high-level brain areas during simple sound retention. We demonstrate the existence of neurons with task-modulated activity in an extensive network of such areas.

We recorded from medial temporal lobe, insula, basal ganglia and cingulate cortex. We hypothesized that neurons in the medial temporal lobe (hippocampus, parahippocampal gyrus and amygdala) would show the highest degree of modulation during the maintenance of tone frequencies. We expected maintenance modulation to be most pronounced in posterior hippocampus, given its specialization for fine-grained information *[11, 12]*. The data demonstrate neuronal responses in high-level cortex with different profiles across task phases and a relationship between the number of neurons modulated during maintenance and working memory performance.

## Results

### Behavior

Ten intracranially-implanted participants completed 60 trials of a non-verbal auditory working memory task (barring one participant who completed 30 trials) where they were instructed to hold a tone in mind and then, after a 3 second delay [“maintenance”], adjust ongoing tones [“adjustment”] to match as closely as possible the frequency of the remembered tone (see **Figure 1** for task schematic and *Methods* for further details). Participants were able to complete the behavioral task to a standard comparable to non-surgical subjects *[13, 14]*, with mean precision estimate scores of 1.04 (± 0.07 standard error) across blocks. These values were not significantly different from estimates derived from *[13]* (*t*(21) = 0.38, *p* = 0.71). To determine whether feedback provided to participants induced learning, we examined performance error across trials (i.e. final distance to target tone during adjustment). Despite receiving this feedback, there was no indication of a significant trend for systematic improvement over time in any participants (individual Mann-Kendall tests, *p* > 0.05 uncorrected for each block). Thus, further analyses involved all trials.

**Figure 1.**
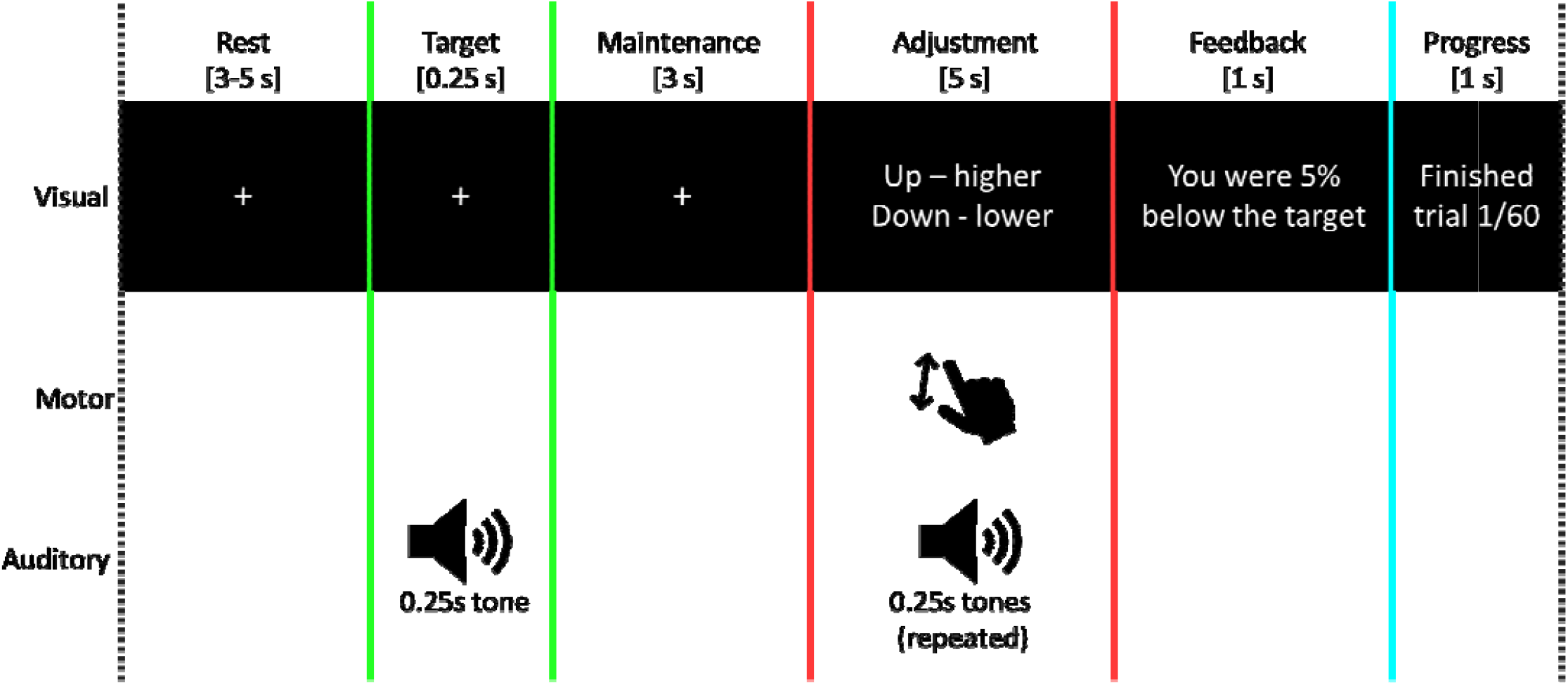
Schematic of the task paradigm. Overview of the task for a single trial. Each participant completed 60 trials (except one who completed 30 trials). Starting tone frequency was randomized between 400 and 1000 Hz for each trial. Colored lines denote task periods that are consistent with those shown in later figures. Note that task phase durations are not to scale.

### Single-neuron data

Participants were implanted with hybrid electrodes *[15]* for the clinical purpose of seizure localization, which allow the isolation of single neurons via microwires that extrude from the tip. **Figure 2a** shows the distribution of electrodes across the brain that yielded single neurons, with regions broadly defined. Across all electrodes, a total of 1269 neurons were isolated (see Supplementary **Figure 1** for curation metrics and **Supplementary Figure 2** for the distribution across brain regions), approximately half of which (∼48%) were located within the medial temporal lobe (MTL).

**Figure 2.**
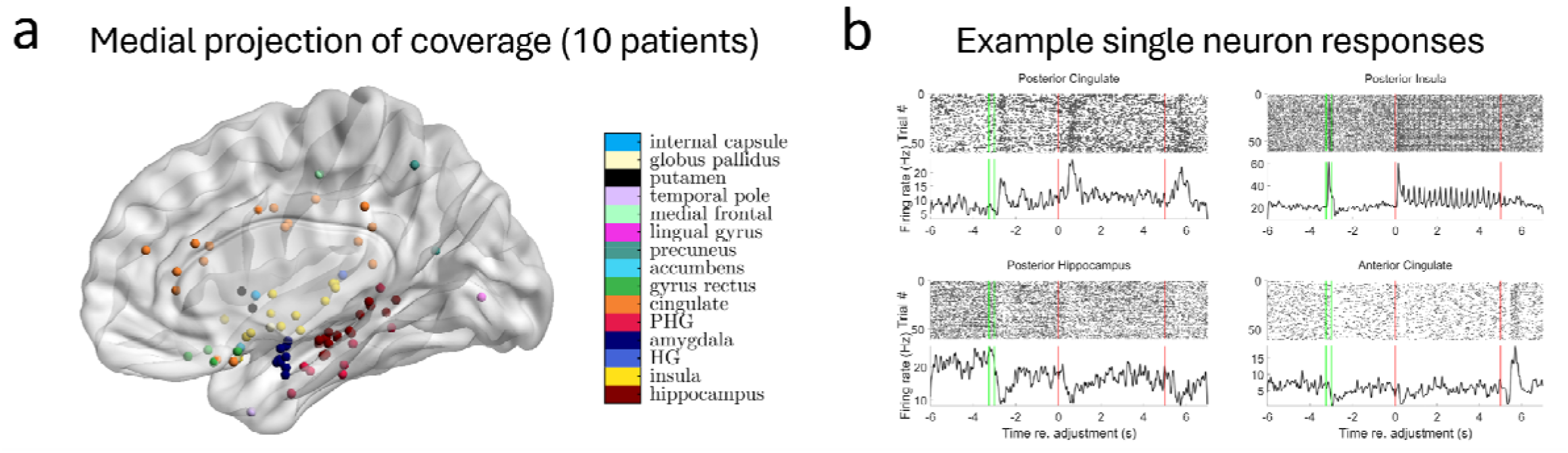
Locations of isolated single neurons and example responses. a, Locations where one or more single neurons were resolved, projected onto a medial view of an MNI152 template reconstruction. b, Example raster plots (top panels) and peri-event spike density functions (bottom panels) for four individual neurons, located in posterior cingulate (top left), posterior hippocampus (bottom left), posterior insula (top right) and anterior cingulate (bottom right). Task phases are denoted by colored vertical bars that are consistent with Figure 1 (maintenance period between second green and first red line; adjustment period between two red lines). PHG: parahippocampal gyrus. HG: Heschl’s gyrus.

Task-related modulation for each neuron was determined for each period of interest based on permutation testing of spike rates across trials compared to baseline values (see *Methods*).

Neuronal dynamics within and between regions were highly variable (see **Figure2b** and **Figure 3**). Some neurons showed increases in activity while a target tone was maintained in memory before a subsequent tone was adjusted to match the target, but many others exhibited patterns of suppression relative to baseline firing rate. This modulation is quantified in **Figure 4** and **Supplementary Table 1**. We performed separate analyses for the first half (-3s to -1.5s prior to adjustment) and second half (-1.5s to 0s prior to adjustment) of the silent maintenance period to assess the persistence of dynamics observed following the target tone.

**Figure 3:**
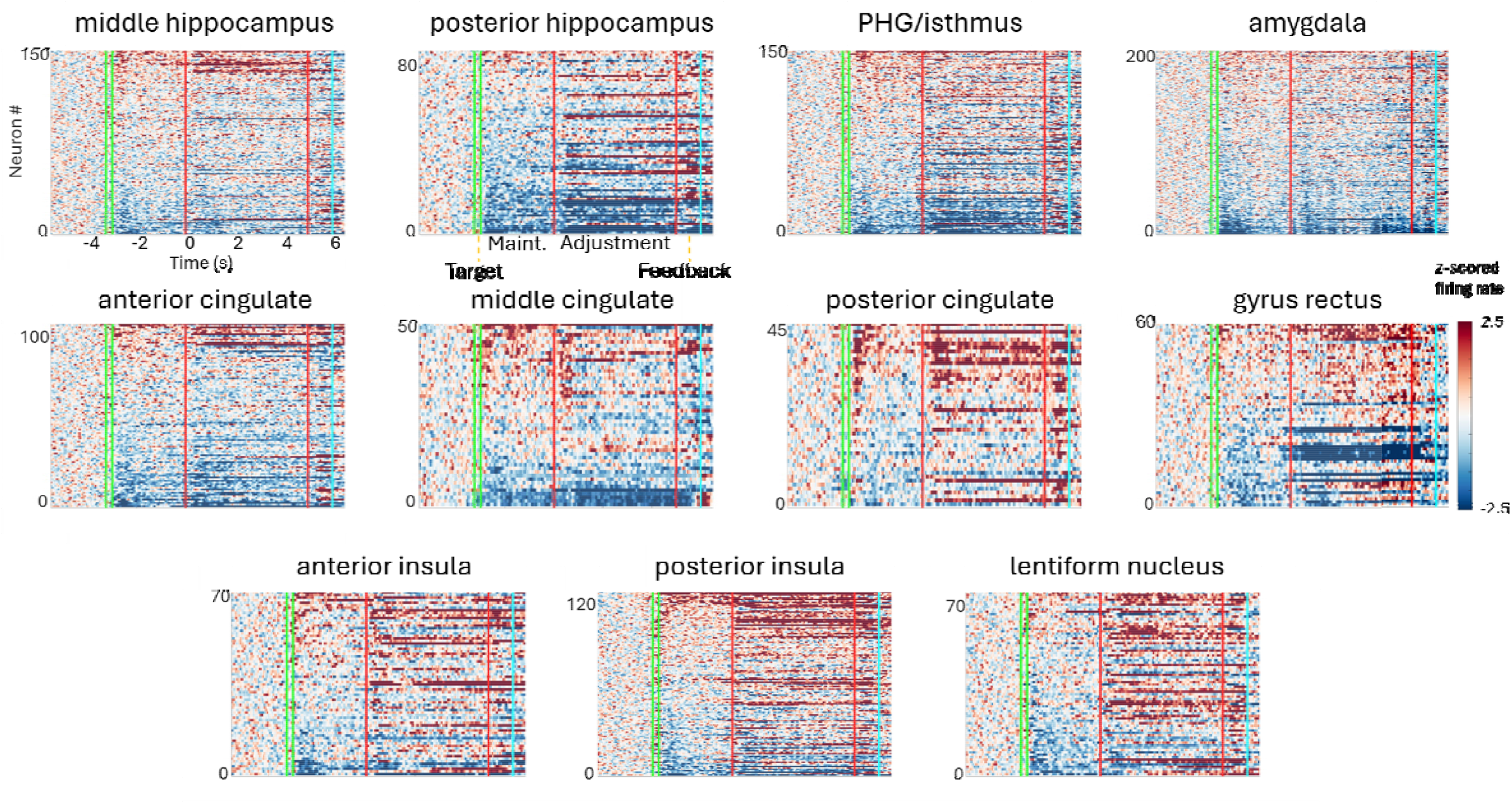
Neuronal responses are heterogeneous within and between regions and task phases. Each panel shows averaged z-scored raster plots for regions with a minimum of 45 neurons isolated. Each row represents the averaged activity of a different neuron, sorted according to the magnitude during the first maintenance period. PHG includes neurons localized to isthmus; anterior cingulate includes neurons within genual and subgenual cingulate. Task phases are specified on second panel and denoted by colored vertical bars. Maint: maintenance period.

**Figure 4:**
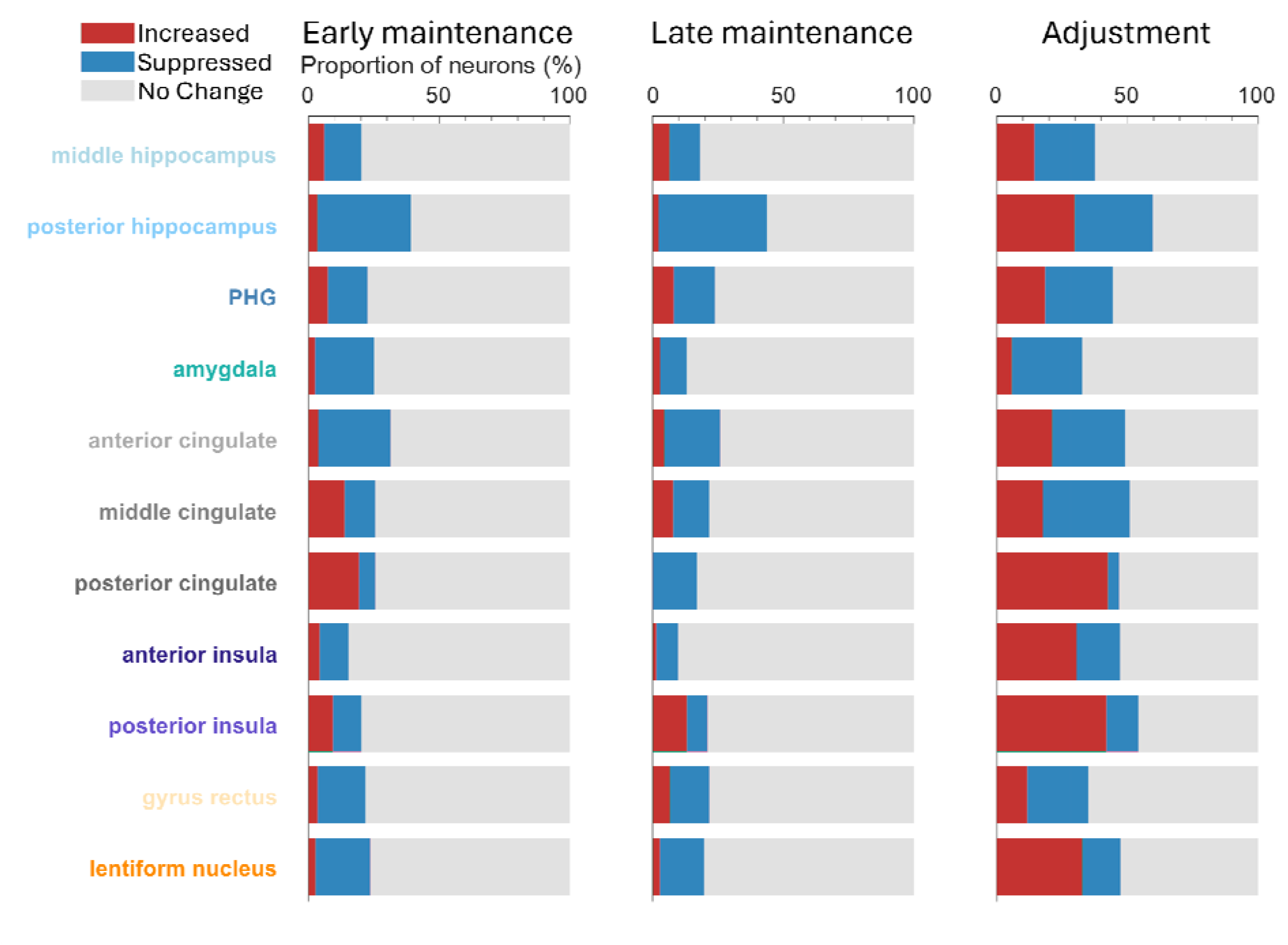
The highest proportion of modulated neurons was in hippocampus with prominent suppression. Proportion of neuronal response types for main task phases of interest, separated according to region from which neurons were recorded and examined across all isolated neurons for a particular region. Early maintenance period is defined as first 1.5 seconds of tone maintenance; late maintenance period indicates 1.5 s to 3 s after tone offset (immediately prior to adjustment phase). Proportion of no change neurons reflects those that did not show significant modulation of firing rates compared to baseline (*p* ≥ 0.05; see *Methods*). Only regions with a minimum of 45 neurons resolved are shown.

The largest proportion of modulated neurons during tone maintenance was in posterior hippocampus at both the start and end of the maintenance period, predominantly showing suppression (∼36% of neurons during first maintenance period, ∼41% during second maintenance period), with only a very small proportion of neurons showing significant increases. During the adjustment period, the proportion of posterior hippocampus neurons exhibiting increased activity was similar to those that decreased (∼30% in each case). This finding contrasts with posterior cingulate, in which a a higher proportion of neurons showed increases in firing rate at the start of the maintenance period, with a lower proportion showing suppression (∼19% vs ∼6%). Of those neurons that were modulated in posterior cingulate during adjustment, the vast majority showed increased firing (∼43% of all recorded neurons). This was also evident in posterior insula (∼42% of neurons increasing during adjustment), which – at least in part – likely reflects the fact that neurons there are responsive to sound, even during passive listening *[16]*.

To avoid biases generated by statistically thresholding each neuron prior to summarizing – for example, missing consistent subthreshold modulation across neurons within a region – we additionally examined the magnitude of changes in firing rates across all neurons for each region (**Supplementary Figure 3**). This clearly demonstrated a significant suppression of MTL neurons during all task phases, although this effect was absent from middle hippocampus and greatest in posterior hippocampus. Anterior cingulate aligned with the MTL in terms of activity patterns at each task phase. Posterior cingulate neurons were significantly suppressed during the second half of the maintenance period and then showed significantly increased activity during adjustment. Lentiform nucleus neurons in the basal ganglia were significantly suppressed at the beginning of the maintenance period and significantly increased during adjustment.

We hypothesized that neuronal states would reach stable plateaus at different phases of the task, based on theories suggesting that attractor states are crucial for working memory *[17, 18]*, with the additional expectation that the population representation would distinguish task phases from each other. We determined the population state-space dynamics using principal component analysis, computing across neuronal spike density functions that were averaged across trials for each neuron and preserving the time dimension. Across all 1269 neurons in all brain areas, 90% of the variance was explained by the first 138 principal components (**Supplementary Figure 4**). The results of this analysis are shown in **Figure 5** with task phases colored separately. Trajectories of neuronal firing traversed distinct regions in state space at different phases of the task. Importantly, these trajectories reached distinct attractor-like states during the maintenance and adjustment phases, suggesting that stable neuronal activity patterns formed during the course of the trial, and that these patterns represented population responses relevant to performing the task. We additionally established the concordance of the timing of these attractor states with the phases of the task using a clustering algorithm (see *Methods* and **Supplementary Figure 5**). Dwell times were observed to divide the set of clusters into longer, attractor-like dwell states and brief transitional states (**Supplementary Figure 5, panel A**). A Gaussian mixture model fit to the distribution of total dwell times determined a threshold dwell time above which clusters were considered to reflect attractor-like states (all-trial threshold: 1.14s; high-error trial threshold: 0.66s; low-error trial threshold: 3.48s).

**Figure 5:**
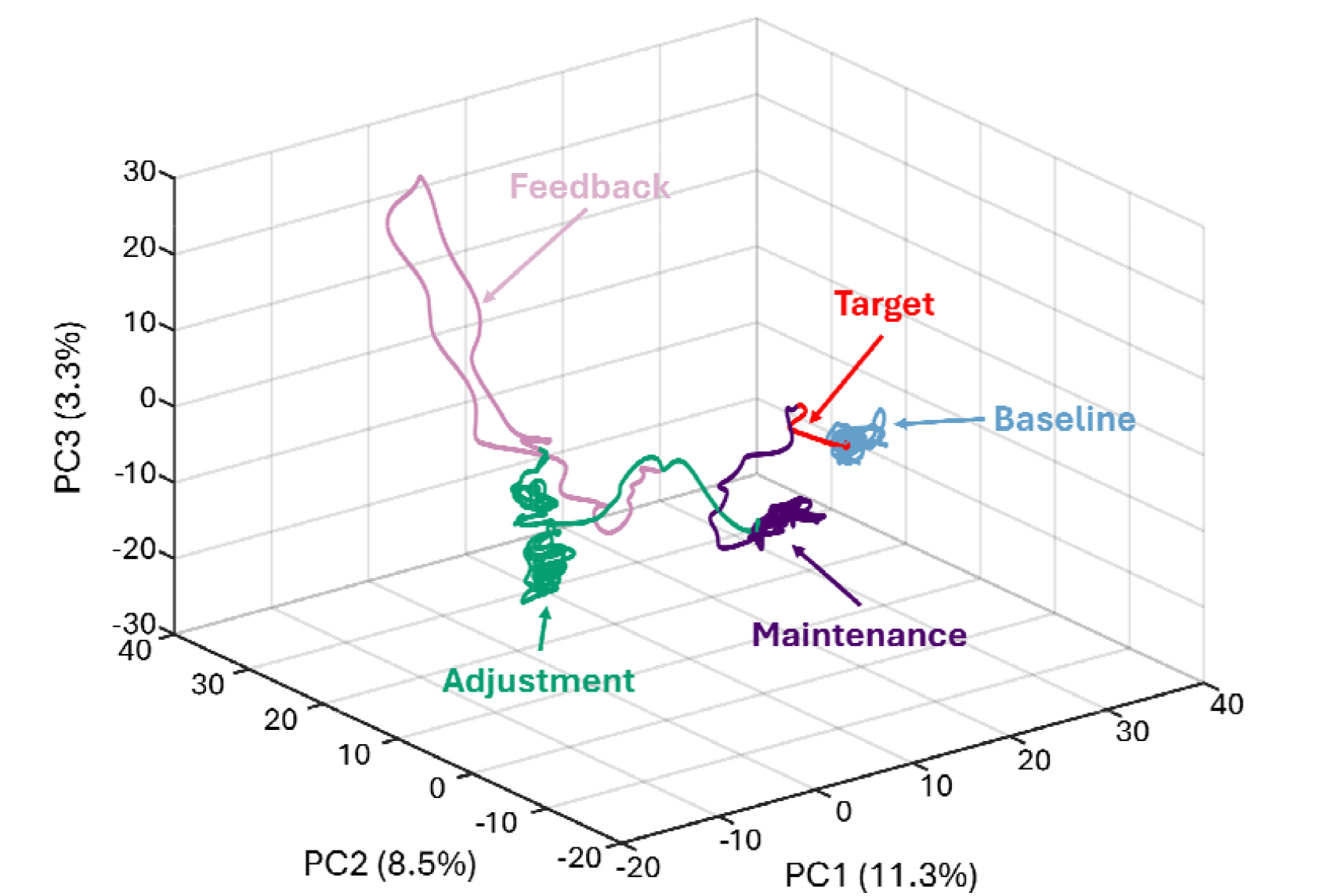
Task phases are clearly differentiated by state space trajectory. State space projection of neuronal trajectory throughout the task across all isolated neurons. First three principal components are shown, with variance explained shown in parentheses for each principal component. The trajectory of all neurons was examined based on spike density functions constructed across all trials for each neuron and z-score normalized prior to principal component analysis. For this figure, the following time definitions were used: Baseline = -5.5 -3.25 s; Target = - 3.25 to -3 s; Maintenance = -3 to 0 s; Adjustment = 0 to 5; Feedback = 5 to 6.5 s.

Within the all-trial PCA trajectory, 3 attractor-like segments were identified (**Supplementary Figure 5, panel B**), each one falling largely within a different phase of the task: one within the baseline phase of the task (segment time range: -5.12s to -3.14s), one within the maintenance phase (-2.87s to 0.19s), and one within the adjustment phase (0.46s to 5.32). Individual trajectory segments were also scored by recurrence rate (RR), which reflects the observed tendency of a given trajectory to return to previously visited locations (see *Methods*). Different attractor-like states exhibited differential recurrence rates (baseline segment: RR = 1.09, maintenance segment: RR = 2.43, adjustment segment: RR = 0.76). We had hypothesized that an attractor-like trajectory within the maintenance phase might reflect a dynamical system for working memory critical to the retention of the stimulus under this task; compared to the trajectories observed during the baseline and adjustment phases, the substantially higher RR of the maintenance trajectory was consistent with this notion. Based on a normalized mutual information (NMI) analysis, which reflects the extent to which knowledge of one random variable reduces the uncertainty of another (see *Methods* for full details), we found a significant alignment between the timing of the detected attractor-like states and corresponding labeled task phases (**Supplementary Figure 5, panel C**, NMI = 0.71, *p* = 0.0414; permutation tested, 20,000 replicates). This constitutes further evidence that the timing of attractor-like states observed within the single-unit-derived PCA trajectories reflects dynamics associated with the phases of the WM task.

We then examined whether neuronal data predicted task performance and whether we could decode the sensory trace from neuronal activity. Comparisons between behavioral and neural data were first made based on decoding analyses by implementing a maximum-correlation-coefficient classifier, following separation of trials into low or high error (see *Methods*). We did not find any reliable decoding of single-trial error (defined by a median split of the final distance from the target frequency across trials for each participant) nor the tone frequency (defined by a median split into low- and high-frequency tones across trials for each participant) during the maintenance period. Decoding of behavioral error briefly reached above chance at the beginning of the adjustment phase (**Supplementary Figure 6**). During the feedback phase, when participants were told how well they had performed on that trial, decoding of behavioral error was significantly above chance, which may suggest performance monitoring.

We also hypothesized that the number of neurons recruited to perform the task may be a predictor of behavioral performance, as in a visual working memory study *[19]* in which the number of modulated maintenance neurons in correct trials was higher than in incorrect trials. Statistical comparisons of neural data were made separately for low and high error trials using permutation testing against baseline as previously described (see *Methods*). As shown in **Figure 6**, across brain regions the number of neurons showing significant modulation was higher during the first 1.5 seconds of the maintenance period when participants performed the task well [Low error trials] relative to when they made more errors [High error trials] (*X*^2^ = 12.97, *p* < 0.001, Bonferroni-corrected), while differences were not significant for the other two task phases. We implemented a generalized linear mixed effects model based on the data in **Figure 3** to determine whether the ability to predict modulation from trial accuracy varied with brain region (see *Methods*). Based on this model, the greatest behaviorally related increase in the number of neurons showing significant modulation was in PHG/isthmus (49% compared with 36% or less elsewhere), however the interaction between region and a trial’s error status was not significant in predicting whether a given neuron showed such modulation. Combined with the decoding results, this suggests that the firing rates did not systematically differ as a function of performance but that a larger number of neurons showing modulation was associated with completing the auditory WM task more successfully.

**Figure 6:**
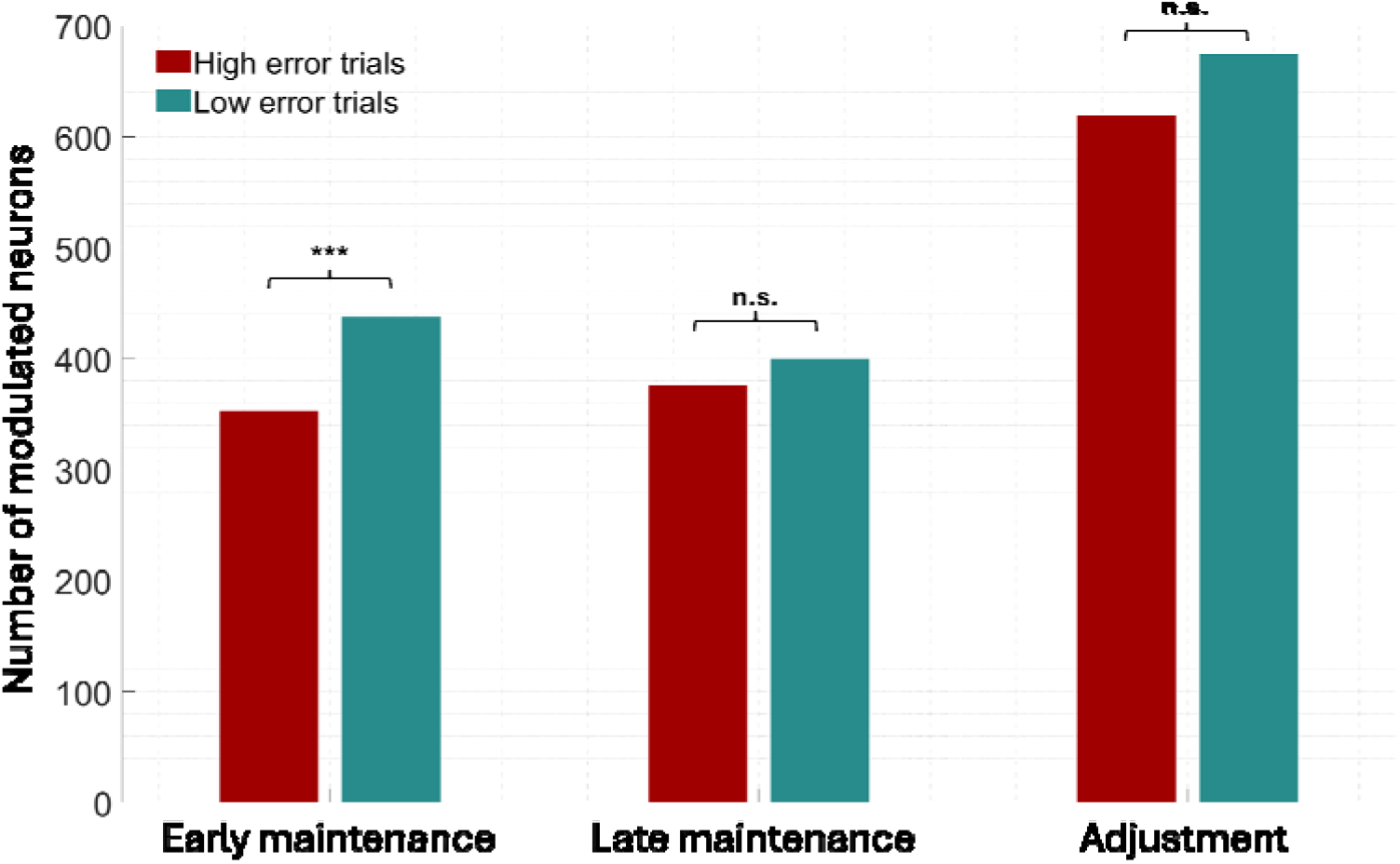
A larger number of neurons were modulated when behavioral performance was better. Number of significantly modulated neurons, separated according to task phase and median split of low (red bars) vs high (green bars) error trials. *** *p* < 0.001, Bonferroni-corrected, n.s. not significant.

Differences between high- and low-error trial responses were also examined through the lens of the per-patient all-unit PCA trajectories, this time segmenting them solely based on the known trial phase timings into *baseline, target, maintenance, adjustment* and *feedback* trajectories. For the 3-dimensional trajectory associated with each task phase, we characterized the differential recurrence rates (RR; see *Methods*) of the segments under each condition. It was hypothesized that the RR of the attractor-like trajectories (reflecting the degree to which a dynamical system revisits prior states and therefore the stability of the attractor) would be elevated under the low-error condition relative to the high-error condition, and particularly within the maintenance trajectory, where a higher RR might reflect a more stable mnemonic representation of the stimulus. Notably, a significant difference was observed between the RR under the low- and high-error conditions during the adjustment phase, at the group level (N = 13 patients, *p* = 0.0017, Wilcoxon signed rank, Bonferroni-corrected α = 0.01), with a higher median RR under the low-error condition than the high-error condition, suggestive of stronger attractor-like recurrence during low-error trials (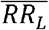 denotes low-error median RR, 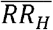 denotes high-error median RR; Adjustment phase: 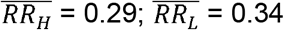). Group-level differences between the RRs of the remaining phases were not significant after multiple comparisons, but demonstrated a consistent effect direction, whereby RR was near-universally higher under the low-error condition (Baseline: 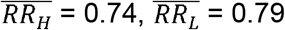, *p* = 0.13; Target: 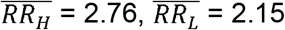, *p* = 0.59; Maintenance: 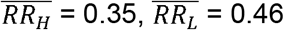, *p* = 0.048; Feedback: 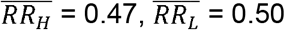, *p* = 0.094), with the exception of the extremely short target phase. This is consistent with our hypothesis that greater attractor stability may facilitate higher performance, and suggests performance on this task may be supported by greater attractor-like activity during the adjustment phase in particular.

## Discussion

For the first time, we examined activity in a large population of human single neurons during a non-verbal auditory working memory paradigm, demonstrating clear modulation within areas beyond auditory cortex. We saw heterogeneity in neuronal responses where individual neurons responded at different phases of the task with either suppression or increase in spiking activity. Between the areas we saw differences in the proportions of these neuronal response patterns. While there was clear modulation in all regions examined, the majority of recorded neurons did not show modulation of activity, suggesting that techniques with poorer spatial resolution may not be sensitive enough to indicate involvement of these regions in auditory working memory.Furthermore, dimensionality reduction applied to the population activity across the brain clearly distinguished different task phases, with striking separation between encoding, maintenance, adjustment and feedback periods. Attractor-like “dwell states” were detected within multiple phases, most prominently during maintenance and adjustment. At the group-level, during the adjustment phase, per-patient dimensionality reduction revealed significantly higher recurrence in the low-versus the high-error condition, consistent with the notion that behavioral performance may be dependent upon the stability of a latent attractor instantiated by the neuronal population. Finally, there was a significant relationship between the number of neurons modulated at the beginning of the maintenance period and the subsequent task performance.State-space analysis was consistent with greater attractor-like activity during the adjustment phase supporting better performance.

An important aspect of the current work is that we were not examining selectivity for acoustic features in our neurons, which has been shown to be a factor in working memory performance based on visual paradigms. For example, previous studies have a that neurons that are involved in the maintenance of visual stimuli are also active during perception, e.g. *[6, 9, 20]*. It was our deliberate intention to avoid complex stimuli with multiple high-level features in this paradigm. This allows interpretation based on working memory for a fundamental sound property independent of any stimulus tagging with labelling *[21]* or feature conjunction *[22]*.

Theoretical accounts of working memory have proposed that dynamical neural systems form stable states – known as attractors – to enable the storage of sensory information over a short period despite a noisy neural system *[17, 18, 23]*. These stable attractors may be represented by a continuum of states (e.g. in lines or rings) or discrete states (e.g. clustered representations). Previous literature in monkeys has provided some support for the idea of stable attractor states *[24, 25]* and our data also support neurons in the brain demonstrating discrete stable attractor-like states during the maintenance and the adjustment phases of our task.

Our results support the idea that hippocampus – in particular the posterior hippocampus implicated in sensory analysis *[11, 12]* – is involved in working memory maintenance. While this idea has been controversial *[26, 27]*, several recent human single neuron studies support the involvement of this area in visual working memory *[4-9]*. Our results further suggest that PHG, amygdala and anterior cingulate act as a consistent unit with hippocampus during sound maintenance, based on their similar modulation patterns. Posterior cingulate, lentiform nucleus and posterior insula were particularly active during the adjustment phase of our task, suggesting their strong involvement in matching ongoing stimuli to a target. The data from the lentiform nucleus provides further support for the involvement of the basal ganglia in focusing attention and gating information relevant for working memory tasks *[28, 29]*.

Based on a maximum-correlation-coefficient classifier (implemented with *[30]*, **Supplementary Figure 6**) applied to firing rate, across neurons we could predict neither behavioral performance nor tone frequency reliably above chance during the maintenance period. Thus, the number of neurons recruited during maintenance was predictive of a greater success of working memory performance, but the patterns of modulation across neurons (i.e. firing rates) did not predict how well someone performed nor reveal the sensory information maintained. The finding of greater neural recruitment during more successful trials may explain a recent report of behaviorally linked BOLD activity in hippocampus during another auditory adjustment task *[31]*. Prior models of working memory suggest that higher order areas may assist with maintaining a representation within sensory cortex, rather than maintaining the representation itself *[32, 33]*, consistent with our being unable to predict whether the tone was high or low frequency, as neural activity (at least in MTL) would be agnostic to low-level sensory features. Our sample size for auditory cortex was small and only obtained from one participant, so we could not draw definitive conclusions about the sensory representation within the maintenance period there, although we were able to significantly decode whether a target tone was high or low frequency during both target presentation and adjustment phase (see **Supplementary Figure 6**). Future studies are therefore needed to understand the contribution of individual neurons in sensory cortex to auditory working memory and the functional relationship between sensory cortex and the high-level areas we examine here. Overall, these data demonstrate a distributed neural code for working memory for a fundamental sound feature that is not explained by semantic or affective association.

## Acknowledgments

We are grateful to Kirill Nourski and Haiming Chen for assistance with recruitment and data collection.

## Author contributions

Conceptualization: JIB, PEG, SK, TDG

Methodology: JIB, AJB, PEG, TDG

Software: JIB, CKK

Formal analysis: JIB, AJB, RMC

Investigation: JIB, PEG, AER, CMG, HK

Data curation: JIB, AJB, TDG

Visualization: JIB, RMC

Resources: MAH

Writing – original draft: JIB

Writing – review & editing: JIB, AJB, PEG, CKK, AER, RMC, CMG, SK, HK, MAH, TDG

Funding acquisition: MAH, TDG

## Data, code, and materials availability

The summary neural data table will be uploaded to Open Science Framework upon publication of this manuscript. Due to confidentiality, raw patient data are not made publicly available. Further anonymized data may be available on request dependent on data transfer agreements and other required collaborative documents.

## Supplementary Materials

## Materials and methods

### Participants

Extracellular single-neuron recordings were obtained from ten adult neurosurgical patients, implanted with electrodes for the clinical purposes of monitoring epileptiform activity prior to potential treatment. Research was conducted under approval of the University of Iowa Institutional Review Board and written informed consent was obtained from all participants prior to data collection, with verbal consent obtained again before each recording session.Recordings were made while subjects were either reclined in a hospital bed or seated in a chair, in a custom-designed dedicated electromagnetically shielded facility within the University of Iowa Epilepsy Monitoring Unit. A total of thirteen recording blocks were included. For the three participants that repeated blocks, these blocks were separated by a minimum of 24 hours.

### Electrodes and recording setup

Stereo electroencephalography (sEEG) depth electrodes (Ad-Tech Medical, Oak Creek, WI) were placed in brain locations based on a clinical need to identify seizure foci *[34]*. Between 4 – 12 of these sEEG electrodes were of a hybrid design *[35]*, that included 8 insulated high-impedance microwires plus 1 uninsulated microwire (39 µm diameter), all of which protruded from the end, prepared with a cut length between 2 to 5 mm that was dependent on the distance of the most distal macro contact to the appropriate brain target. Each microwire bundle was separated in a splay pattern immediately prior to implantation. These hybrid electrodes therefore enabled the recording of single neurons. Electrode locations were confirmed based on post-operative MRI scans, preprocessed using Freesurfer *[36]*. All neurophysiological data were recorded using a Neuralynx ATLAS System (Neuralynx. Bozeman, MT). High impedance recordings were passed through a preamplifier located on top of the patient’s head (ATLAS-HS-36-CHET-A9, Neuralynx, Bozeman, MT) prior to interfacing with the ATLAS acquisition system. These were subsequently recorded with a 32000 Hz sampling rate, filtered between 0.1 – 8000 Hz and referenced online to the uninsulated microwire.

### Imaging

A T1-weighted structural MRI scan of the brain was conducted for each participant both before and after electrode implantation. The images were captured using a 3T Siemens TIM Trio scanner and a 12-channel head coil. MPRAGE images had a spatial resolution of 0.78 × 0.78 mm, a slice thickness of 1.0 mm, and utilized a repetition time (TR) of 2.530 seconds and an echo time (TE) of 3.520 milliseconds. To locate the recording contacts’ positions on the preoperative structural MRI scans, these images were aligned with post-implantation structural MRIs. This alignment was achieved using a 3D linear registration algorithm (Functional MRI of the Brain Linear Image Registration Tool; *[37]*) and custom-written Matlab scripts (MathWorks, Natick, MA). All recording contacts included here were verified to be within respective locations listed in **Figure 2**. Electrode locations were co-registered for each participant to a template brain, in order to derive MNI coordinates for the purposes of visualization. Contact locations were then visualized across subjects on an ICBM152 template using BrainNet Viewer *[38]*.

### Stimuli and procedure

Auditory stimuli were delivered either diotically using Etymotic ER4B earphones with custom-fit earmolds or via free-field speakers positioned ∼1 m from each ear whenever the earphones were not comfortable for a particular participant. Sound levels were set at a comfortable listening level for each participant and were fixed throughout the recording block. The overall trial structure is shown in **Figure 1**, which was based around an adaptation of a previously published paradigm *[13]*.

For each trial, participants rested for a period randomized between 3-5 seconds whilst watching a monitor screen [“rest”]. They were then presented with a 0.25 second duration tone, with a frequency set between 400 and 1000 Hz and randomized frequency for each trial [“target”].

Following this, there was a 3-second delay wherein they were instructed prior to the task to keep the target tone in mind [“maintenance”]. For all these first three periods, a fixation cross was shown continuously on the screen. Participants were then given 5 seconds to use a keyboard (up or down arrow keys) to adjust repeating tones (0.25-second duration) until the frequency matched as close to the target as they felt they could get [“adjustment”]. The starting tone frequency for this period was randomized on each trial to differ from the frequency of the target tone by between 2 and 10%. All tones were enveloped with 2.5ms onset/offset cosine ramps to avoid transients. Processing time for sound presentation and keyboard input meant that there was ∼5 ms ISI between each tone presentation, which added ∼100 ms to this section of the task. Following this, participants were presented with visual feedback for 1 second informing them how closely they matched the tone [“Feedback”]. Each trial ended with a visual presentation of where they were within the overall task [“Progress update”]. After 30 trials, participants were instructed to take a break for as long as they needed to, and self-initiated continuation of the task. The task lasted approximately 15 minutes in total for each participant to complete 60 trials. One participant decided to cease participation at the break due to tiredness, thus completing 30 trials. The task paradigm was programmed and presented using custom-written MATLAB scripts (MathWorks, Natick, MA) utilizing Psychtoolbox functions *[39]*, and audio was delivered from the computer via a Focusrite Scarlett 8i6 USB interface. Auditory stimuli were aligned with neural data via the sending of event triggers over a parallel port.

### Data processing and analysis

Data from high impedance electrodes were first extracted using Matlab and denoised using the demodulated band transform *[40]*. These data were downsampled to 12 kHz and common average re-referenced to all high impedance contacts on the same assembly prior to spike sorting. Spike sorting was performed using an automated procedure, utilizing a recently developed algorithm *[41]*, and manually inspected for quality purposes. Briefly, filters were estimated for potential candidate single-neuron waveforms for each channel by using higher-order spectral decomposition *[42]*. Extracted features were clustered using a gaussian mixture model in Matlab (R2022a, Mathworks Inc) and spike times from these clustered features were used to plot separate candidate waveforms. Single neurons were defined based on uniformity of waveforms across different spike times for each cluster, along with interspike interval distributions that did not generally violate refractory periods (< 1% of interspike intervals occurring within 1ms). Putative single-neuron spike times were then epoched around for each trial (-6 seconds before adjustment period onset to 7 seconds post-adjustment onset). Raster plots were created to show neuronal activity for each trial and spike density functions were estimated by convolving neuronal spike times with a gaussian kernel (25 ms standard deviation, 1ms resolution). Decoding analyses were implemented with a maximum-correlation-coefficient classifier using the Neural Decoding Toolbox in Matlab *[30]*, based on individual trial firing rates [spike density functions] binned into 200 ms windows, with a 50 ms step size, z-score normalization, 10 cross-validation splits and 20 resample runs.

### Statistical analysis

For assessment of working memory precision, performance on each trial was calculated as the error in semitones between the target tone and the final adjusted tone, as follows,

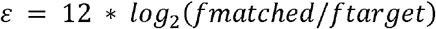

where *fmatched* represents the frequency of the final tone that participants adjusted to on each trial and *ftarget* is the frequency of the initial target tone. Data were median split for each participant to divide trials into low error (i.e. better performance) and high error (i.e. worse performance) for the purposes of relating neural data to behavioral performance. Additionally, working memory precision across trials was estimated by calculating the reciprocal of the standard deviation of the error across trials (*P* = 1/σ), in a similar manner to previous auditory and visual studies *[13, 43]*. This overall precision across participants was compared to previously published data to show similar performance to non-surgical participants *[13]*, based on estimation of mean and standard error values calculated from **Figure 2A** (Memory Load 1) in Kumar et al (2013).

Single-neuron responses during the task phases of interest (early maintenance: -3 to -1.5 seconds prior to adjustment; late maintenance: -1.5 to 0 seconds prior to adjustment; adjustment: 0 – 5 seconds post adjustment onset) were considered significant based on permutation testing against a pre-task baseline period (-5.95 to -4 seconds prior to adjustment onset), with an alpha set at 0.05. A generalized linear mixed effects model was also implemented to examine the relationship between trial accuracy and modulation by region, with a random intercept for recording block, in the form:

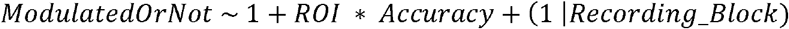

### Recurrence quantification analyses

Principal component trajectories were derived from trial-averaged spike density functions across units (yielding a single samples × 3 trajectory per condition). To extract attractor-like dwell states from these 3-dimensional trajectories, points were first subjected to “density-based spatial clustering of applications with noise” (DBSCAN) clustering [44], yielding distinct clusters of points within the trajectory. DBSCAN is an iterative algorithm that partitions data into both separable clusters and noise via density-based clustering, gradually forming labeled clusters comprising *core points* (which, once labeled, are typically not relabeled) and *border points* (which are typically labeled initially as noise but may later be relabeled as belonging to an existing cluster). To form clusters, DBSCAN relies on two tunable parameters: *ε*_*DB*_, a search radius defining the neighborhood of a given point; and *k*, the minimum number of points required within the neighborhood for a point to be labeled as *core*. Here, *k* was fixed at 60 (20 times 3 dimensions), whereas *ε*_*DB*_ was defined as the value at the knee of the sorted (*k*+1)-th nearest neighbor distances computed on the full trajectory. Onset/offset times and recurrence rate (RR) were then computed for each cluster. Recurrence rate reflects the density of recurrent events within a trajectory. Here, as is common, RR was defined as the proportion of all pairs of observed points wherein the two points were closer than a given radius, epsilon (*ε*_*RR*_)[45], retaining only the upper triangle of the distance matrix, and excluding points less than or equal to *w*= 10 samples apart. The value *ε*_*RR*_ was defined as the 2^nd^ percentile of a defined subset of all pairwise distances, which comprised the upper triangle of the distance matrix computed between all points, excluding those points less than or equal to *w* samples apart. The 10- sample distance constraint was imposed to reduce bias introduced by the inclusion of temporally adjacent samples within the immediately local trajectory.

The number of samples within each detected spatial cluster followed a bimodal distribution of high and low count clusters, broadly correspondent to longer, attractor-like (AL) dwell states and brief transitional states. A Gaussian mixture model comprising two distinct probability density functions (one long dwell and one short dwell distribution) was therefore fit to the distribution of sample counts, with the threshold sample count for AL states defined as the point of intersection between the two distributions. Only clusters with sample counts strictly greater than this cutoff were retained as AL trajectories during automatic clustering.

For each AL trajectory, onset and offset times were ascertained and compared to the known times of each task phase (see above) using normalized mutual information scoring (*MI*_*norm*_; see [46]) between the vectors of ground truth (*g*) and estimated (*c*) task-phase labels at each sample index, computed thus:

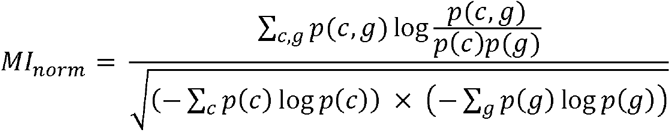

To compute the significance of the observed score, permutation testing was performed (20 000 replicates) in which the originally observed cluster labels *c* were circularly shifted by a random number of samples. Permuting by circularly shifting maintained the temporal dependence between samples within *c*_*replicate*_ (likewise constraining the observed vector). Preceding permutation, the original cluster labels *c* were generated by performing one-to-one remapping of the DBSCAN output such that each label (excluding 0) was reassigned according to which known task phase it maximally occupied, mapping every label in order of most to least prevalent, and remapping if a new label better overlapped with an existing phase than a previous choice. The maximum overlap remapping procedure was performed on the assumption that each attractor-like state in *c* was likely to pertain to the task phase it maximally occupied. To be conservative, this mapping was held constant during permutation.

To conduct between-condition comparisons of the RR within specific segments of the per-patient PCA trajectories, clusters were not determined automatically but instead defined using the known task phase onset and offset times, yielding 5 distinct trajectories per condition per patient. As with the full-cohort PCA trajectories, per-patient PCA trajectories (those constructed from only the single unit responses of a given patient) were derived from trial-averaged spike density functions (SDFs). Wherever in the study RR was reported by condition (i.e. when computing both the descriptive statistics on all-cohort PCA trajectories, and the inferential statistics on per-patient PCA trajectories), *ε*_*RR*_ was first estimated on an all-trial trajectory and held constant for condition-specific analyses, to ensure RR was comparable between conditions. To ensure PCA trajectories were compared within a common coordinate system across conditions, a set of common PCA coefficients (column vector *w*_*A*_) and variable means (*μ*_*A*_) were first computed on the samples × units matrix of all-trial mean SDFs, *X*_*A*_ . High- and low-error condition mean SDFs (*x*_*high*_ and *x*_*low*_, respectively) were then projected to the common space via the following standard projection formula:

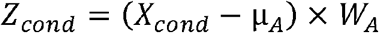

Finally, to test the null hypothesis that, across patients, the per-cluster differences in RR between conditions were drawn from a distribution with a median of 0 (i.e., in order to reject the null hypothesis of no effect), the Wilcoxon signed rank test was applied to the RR condition contrasts for all patients within a given cluster (with α = 0.05 before correction for multiple comparisons). To correct for multiple comparisons, α was Bonferroni-corrected (0.05 / 5 clusters = 0.01).

**Supplementary Table 1.** Proportions of modulated neurons for different task phases. PHG: parahippocampal gyrus.

| Proportion of neurons<br>Region | Early maintenance |  |  | Late maintenance |  |  | Adjustment |  |  |
| --- | --- | --- | --- | --- | --- | --- | --- | --- | --- |
|  | ↑ | ↓ | — | ↑ | ↓ | — | ↑ | ↓ | — |
| middle hippocampus | 5.8 | 14.3 | 79.9 | 6.5 | 11.7 | 81.8 | 14.3 | 23.4 | 62.3 |
| posterior hippocampus | 3.4 | 35.6 | 60.9 | 2.3 | 41.4 | 56.3 | 29.9 | 29.9 | 40.2 |
| PHG | 7.3 | 15.2 | 77.5 | 7.9 | 15.9 | 76.2 | 18.5 | 25.8 | 55.6 |
| amygdala | 2.4 | 22.6 | 75.0 | 2.9 | 10.1 | 87.0 | 5.8 | 26.9 | 67.3 |
| anterior cingulate | 3.7 | 27.8 | 68.5 | 4.6 | 21.3 | 74.1 | 21.3 | 27.8 | 50.9 |
| middle cingulate | 13.7 | 11.8 | 74.5 | 7.8 | 13.7 | 78.4 | 17.6 | 33.3 | 49.0 |
| posterior cingulate | 19.1 | 6.4 | 74.5 | 0.0 | 17.0 | 83.0 | 42.6 | 4.3 | 53.2 |
| anterior insula | 4.2 | 11.1 | 84.7 | 1.4 | 8.3 | 90.3 | 30.6 | 16.7 | 52.8 |
| posterior insula | 9.3 | 10.9 | 79.8 | 13.2 | 7.8 | 79.1 | 41.9 | 12.4 | 45.7 |
| gyrus rectus | 3.3 | 18.3 | 78.3 | 6.7 | 15.0 | 78.3 | 11.7 | 23.3 | 65.0 |
| lentiform nucleus | 2.6 | 21.1 | 76.3 | 2.6 | 17.1 | 80.3 | 32.9 | 14.5 | 52.6 |

**Supplementary Figure 1.**
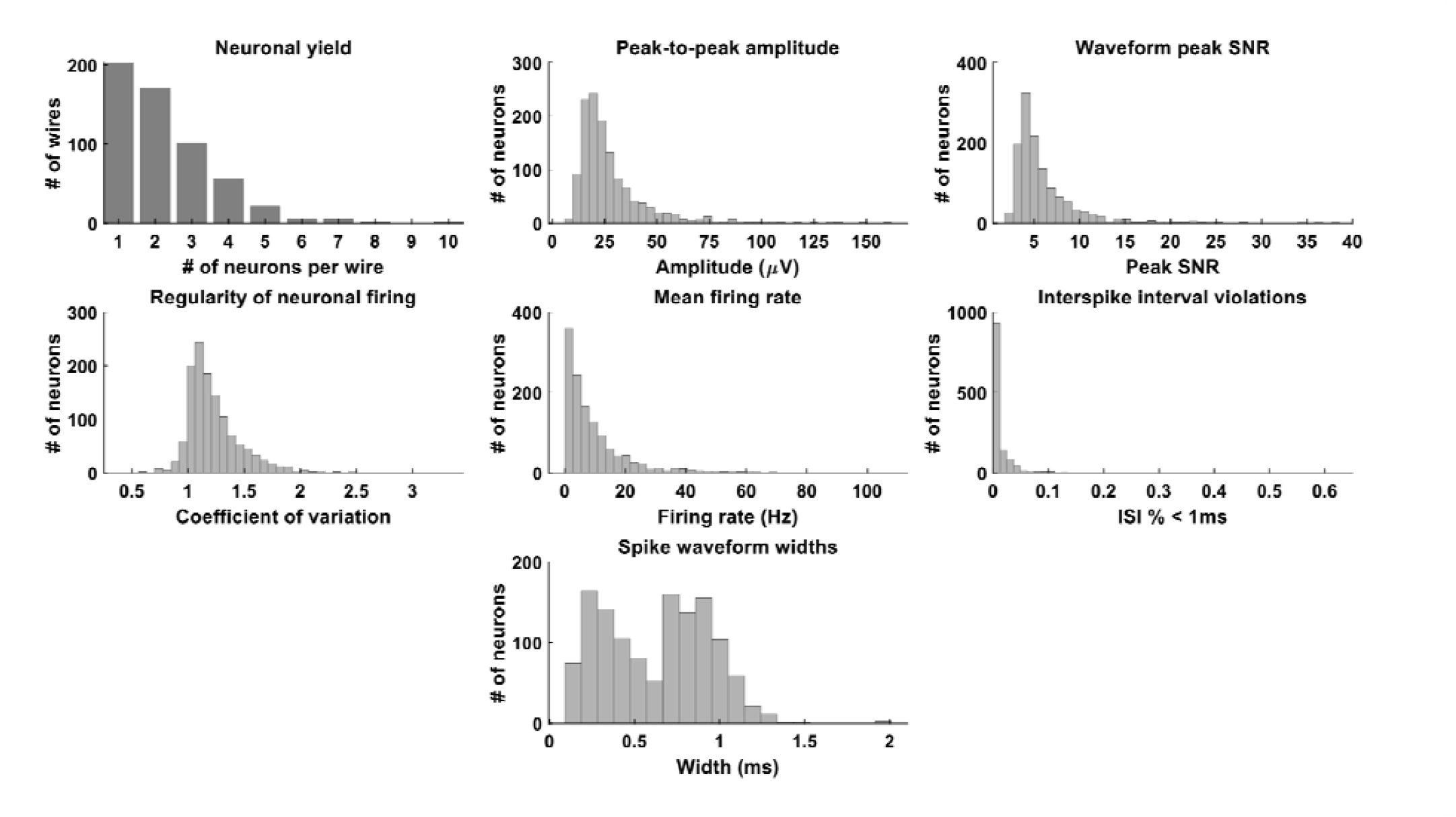
Curation details for all neurons. ISI: inter-spike interval. SNR: signal-to-noise ratio, calculated as the ratio of the peak of each single neuron waveform divided by the median absolute value of the signal for that channel. The bimodal nature of spike waveforms widths as shown in the bottom panel has also previously been observed by others, e.g. *[47]*

**Supplementary Figure 2.**
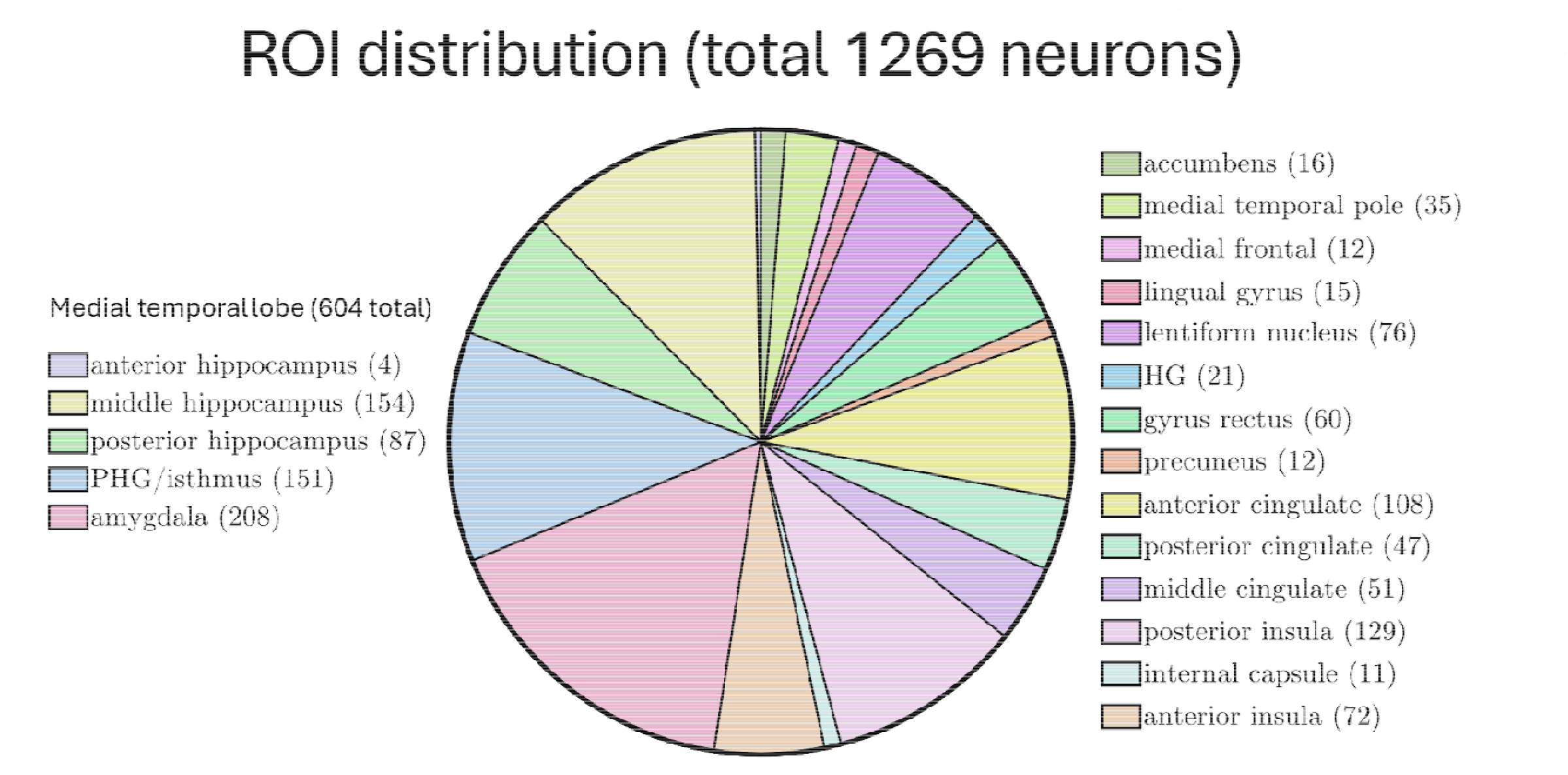
Pie chart showing distribution of locations of all single neurons. Number of neurons resolved is shown in parentheses for each location. Naming is shown from top to bottom for the relevant half of the chart. Lentiform nucleus encompasses globus pallidus and putamen. Almost half of all resolved neurons were located within medial temporal lobe.PHG: parahippocampal gyrus; HG: Heschl’s gyrus.

**Supplementary Figure 3.**
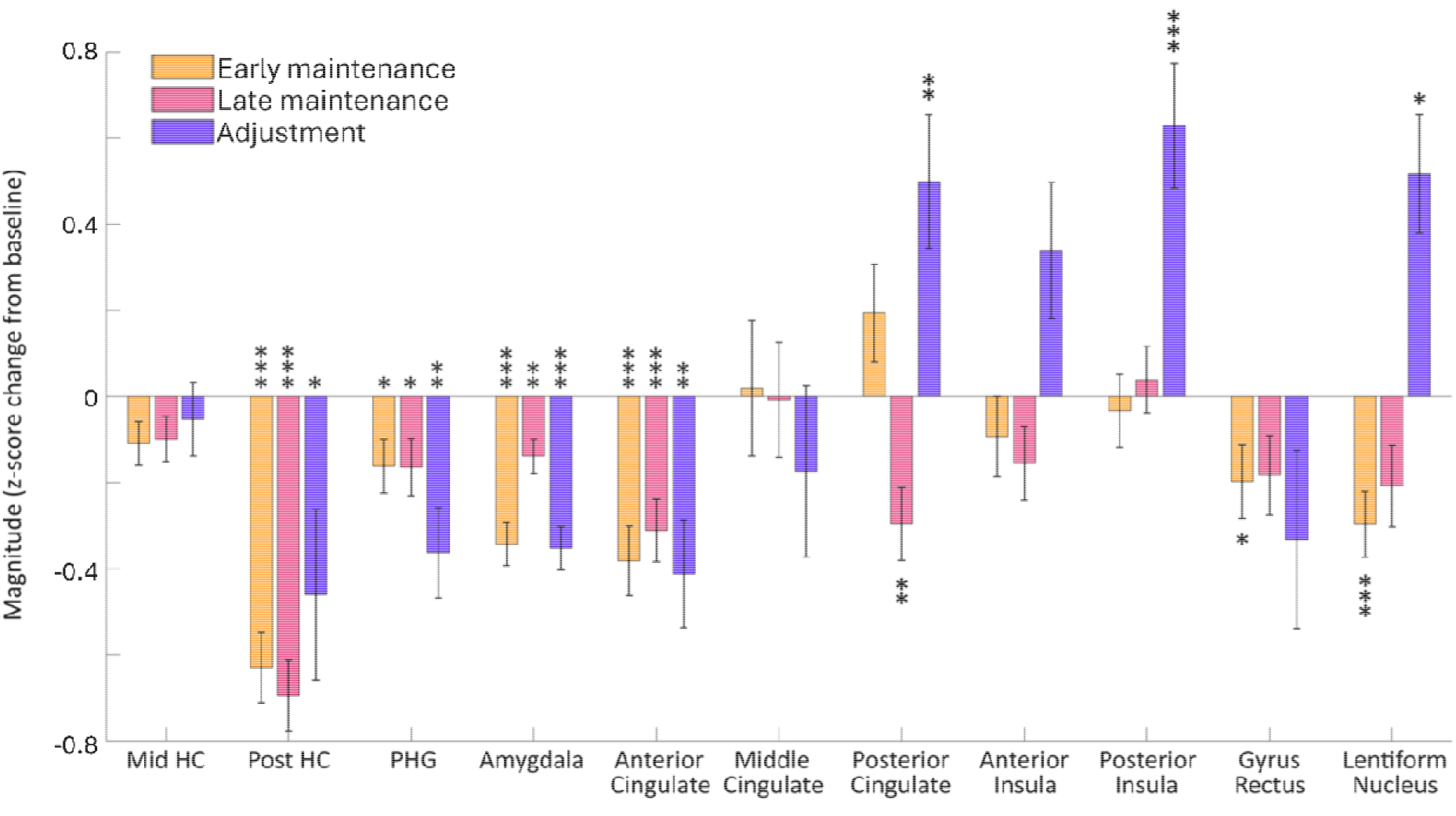
Magnitude of changes at task phases of interest. These were calculated for each neuron based on *z*-score relative to baseline (see Methods), then averaged across all isolated neurons within a region. Error bars represent standard error of the mean. Stars indicate significance of deviation from zero, derived from *p*-values based on one-sample t-tests with False Discovery Rate correction applied (* *p*< 0.05, ** *p*< 0.01, *** *p*< 0.001).

**Supplementary Figure 4.**
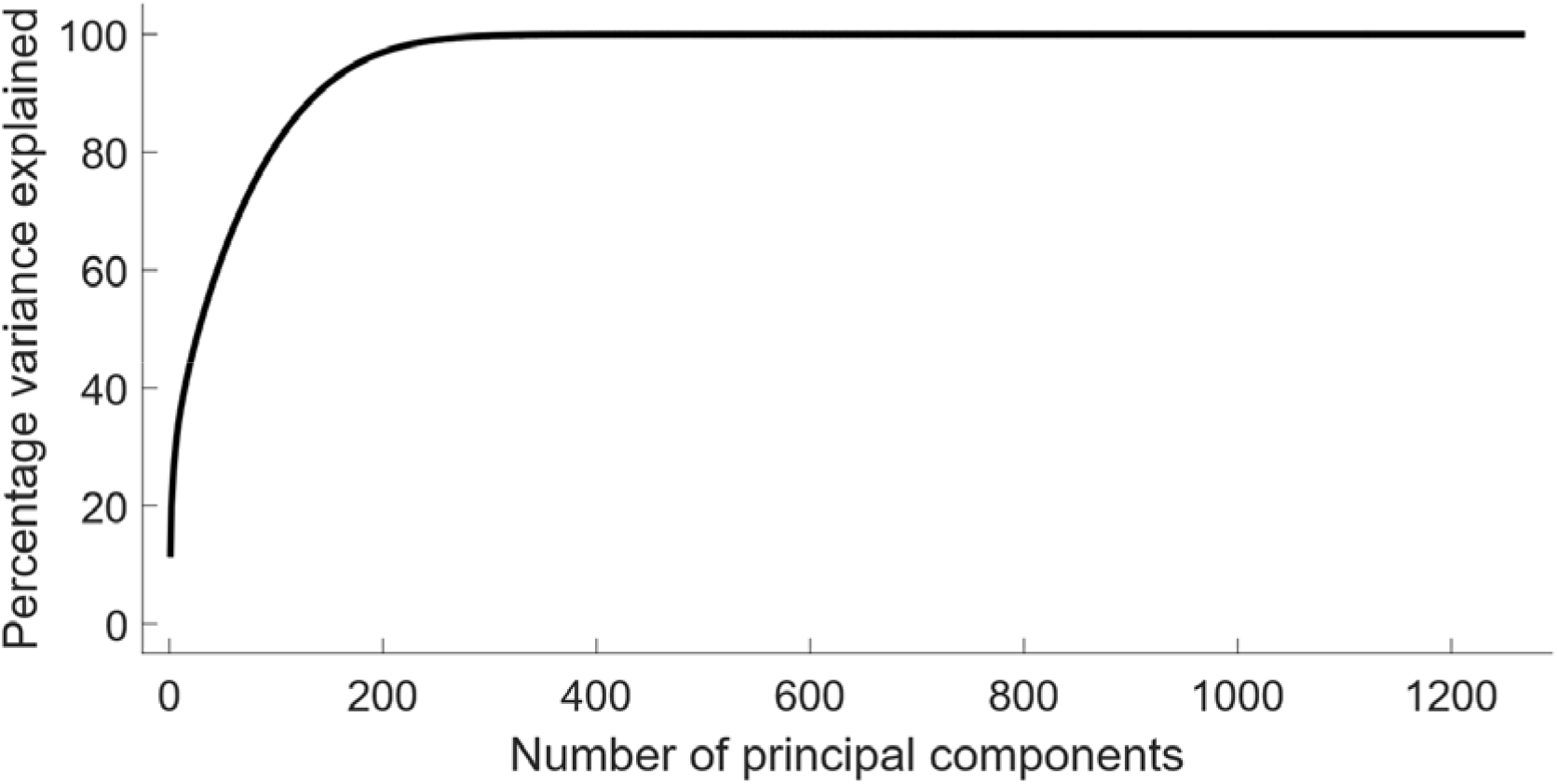
Cumulative variance explained as a function of the number of principal components across all isolated neurons.

**Supplementary Figure 5.**
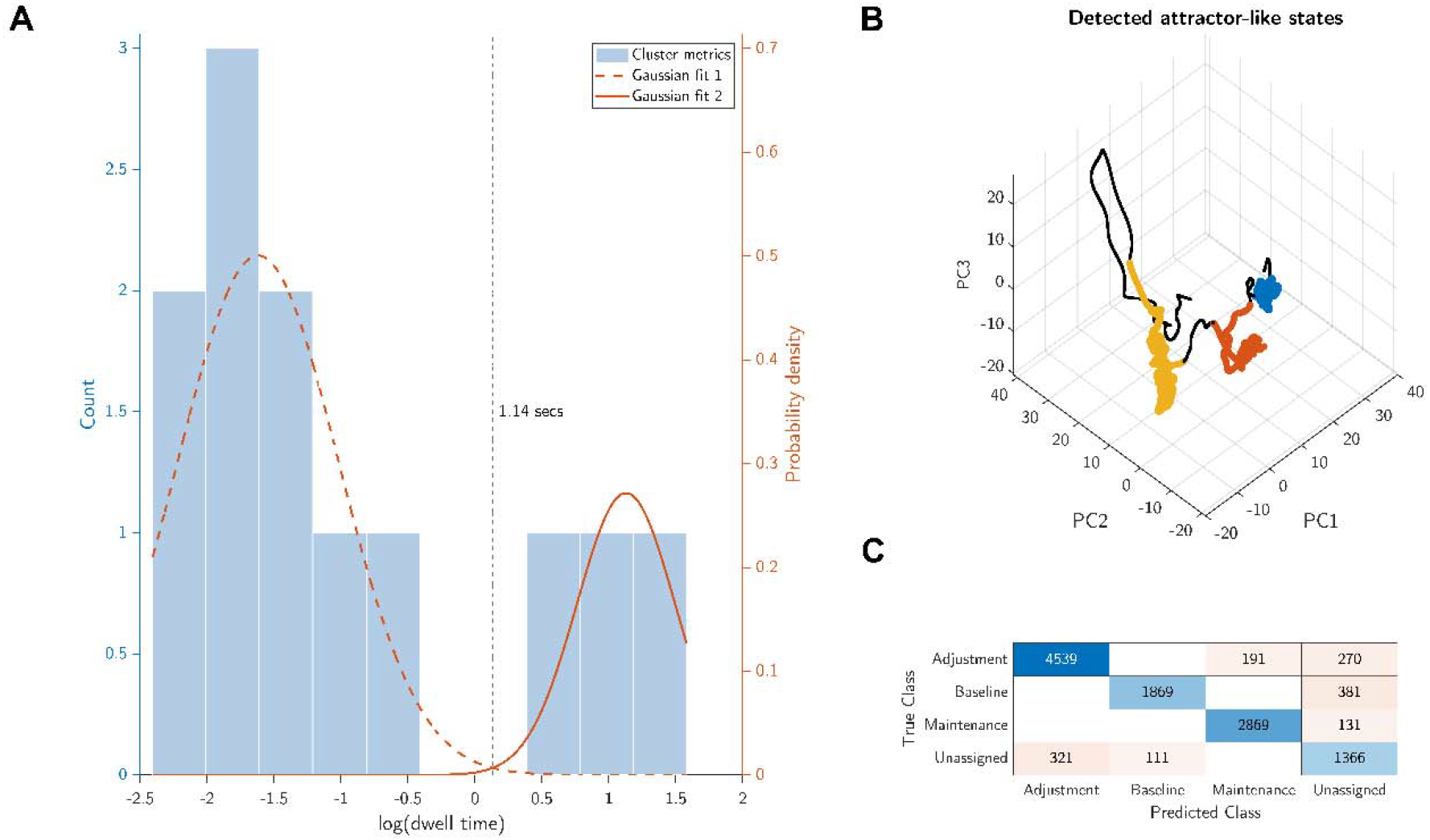
**A:** Histogram of computed dwell times across all DBSCAN-detected clusters. Gaussian mixture model outputs are shown (red solid and dashed lines), revealing distinct categories of high and low dwell time clusters. Vertical black dashed line indicates the intersection point of the two Gaussian distributions. **B:** State space projection of neuronal trajectory throughout the task across all isolated neurons, colored here according to highlight the three detected attractor-like states (dwell time > 1.14 secs), with blue substantially overlapping the baseline phase, red substantially overlapping the maintenance phase and yellow substantially overlapping the adjustment phase of the task. **C:** Confusion matrix showing the correspondence between the predicted class of each sample point based on the attractor-like cluster labels and the ground truth task phases.

**Supplementary Figure 6.**
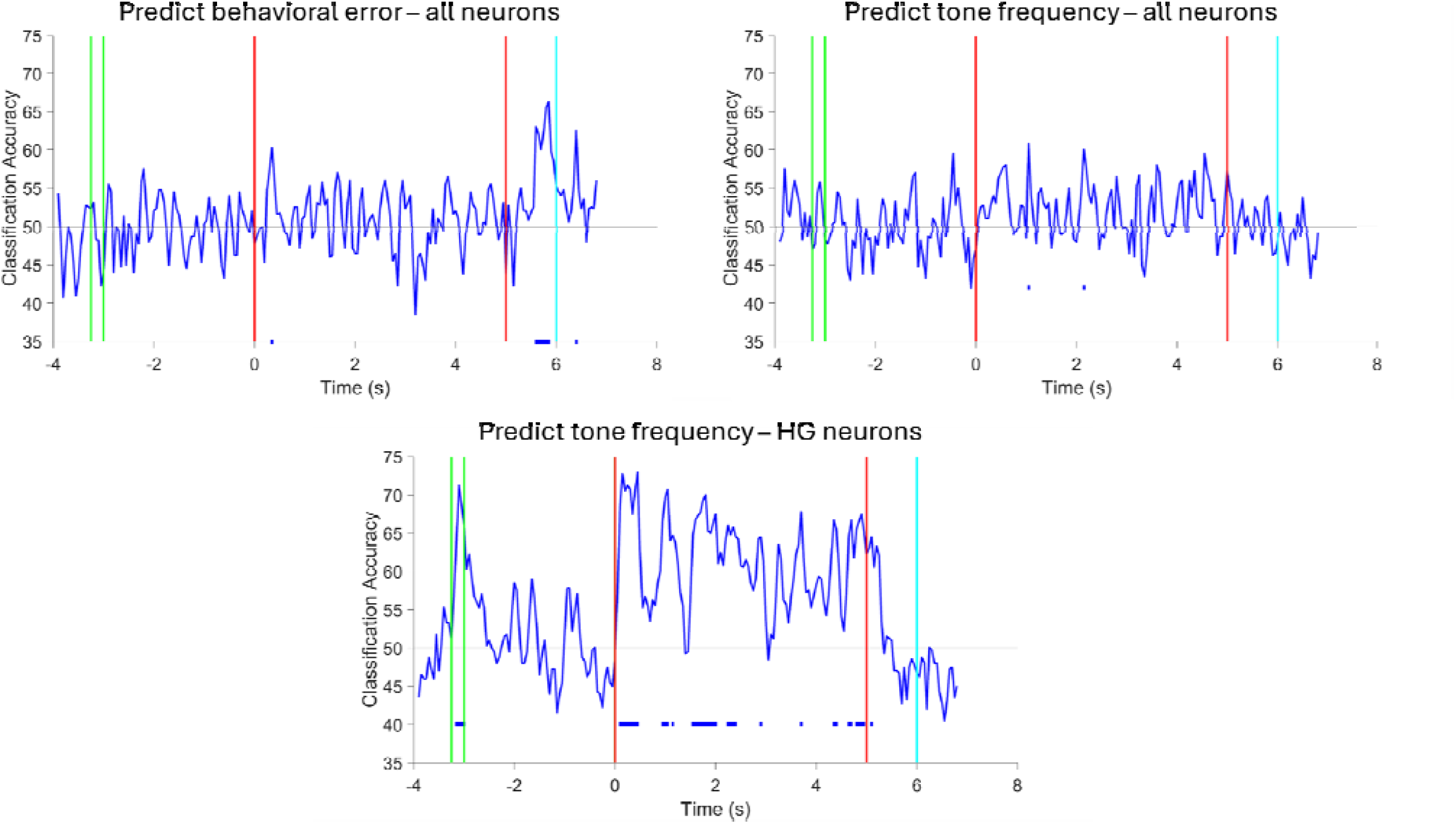
Decoding analyses to determine behavioral error across all neurons (upper left panel), tone frequency across all neurons (upper right panel) and tone frequency for Heschl’s gyrus (HG) neurons (bottom panel), using a maximum-correlation-coefficient classifier. The latter analysis was mainly included to demonstrate feasibility of the decoding approach, as the sample size was limited for this region (21 neurons in one patient). Periods of significance shown with blue horizontal lines below decoding traces, determined via comparison of true label decoding performance against shuffled trial labels to create a null distribution (shuffled 500 times, alpha value set at 0.002, i.e. 1/500). Horizontal black line indicates chance performance. It is important to note that we constrained the starting tone frequency during adjustment to between 2% and 10% of the target frequency, which highlights that adjustment tones began within a similar frequency range to target. Across all neurons, decoding performance was not significantly above chance at any time during maintenance, for either behavioral error or tone frequency. Behavioral error was significantly above chance for a prolonged window during feedback, suggestive of performance monitoring, and briefly above chance at one 50ms window near the start of adjustment, while tone frequency was only briefly above chance during two separate 50 ms windows during adjustment when considering all neurons.

